# Crimean-Congo Hemorrhagic fever (CCHF) micro-neutralization assay using transcriptionally competent virus-like particles (tc-VLPs) in BSL-2 lab

**DOI:** 10.64898/2026.11.25.769095

**Authors:** Stephanie Devignot, Ali Mirazimi, Friedemann Weber

## Abstract

Crimean-Congo hemorrhagic fever (CCHF) virus infection can result in a severe and deadly disease in humans. There are currently no specific countermeasures, so the virus has to be handled in a high-containment, Biosafety Level 4 (BSL-4) laboratory. This is limiting the research and some diagnostic assays to a few labs in the world.

We took advantage of our transcriptionally-competent virus-like particle system (tc-VLPs) for CCHFV to develop and optimize a micro-neutralization assay that can be used in BSL-2 conditions. The tc-VLPs express the structural proteins of the virus, but lack the viral genome: They are unable to produce new particles on their own, and can be safely used under BSL-2 conditions. Moreover, activity of the tc-VLPs can be detected by a luciferase assay, as they express a CCHFV-promoter-driven nanoluciferase reporter gene.

Here, we describe the 3-day procedure to perform a CCHF tc-VLP micro-neutralization assay in a 96-well plate format. It is able to detect neutralizing antibodies in sera and in libraries of purified monoclonal antibodies, and to obtain a neutralization titer (NT50). This combination of a safe BSL-2 assay with a simple luminescence read-out may allow more laboratories to assess the presence of neutralizing antibodies in samples.

Key words: Crimean-Congo hemorrhagic fever virus, CCHFV, Virus-like particles, VLP, Transfection, Neutralization, BSL-2

## 1 Introduction

Crimean-Congo hemorrhagic fever virus (CCHFV, systematic name *Orthonairovirus Haemorrhagiae*) is a member of the *Nairoviridae* family that is transmitted by Hyalomma ticks and by contact with infectious fluids. Infection can result in a severe and deadly disease in humans, with a mortality rate up to 40% ***(1, 2)***. It is the most widespread hemorrhagic fever virus in the world, and there is currently no specific treatment and no approved vaccine. Because of the lack of medical countermeasures, the virus has to be handled in a high-containment laboratory. There are few Biosafety Level 4 (BSL-4) laboratories in the world, and their access is highly regimented and limited, impeding the research on the disease and the virus, and limiting the tools for diagnostics.

Transcription-competent virus-like particles (tc-VLPs) are a powerful and versatile tool: they express all the structural proteins of the virus, including the highly antigenic surface proteins, but lack the viral genome. Instead, they contain a virus-specific minigenome, encoding a reporter gene under the control of a viral promoter. They are hence unable to replicate and to make new particles. As tc-VLPs are non-infectious, they can be handled in Biosafety level 1 or 2 (BSL-1, BSL-2) laboratories, which are available worldwide.

We had developed a CCHF tc-VLP system, which produces particles that are similar to the native virus: The tc-VLPs bear the antigenic glycoproteins Gn and Gc, and contain a CCHFV-specific-promoter nanoluciferase-reporter minigenome that is bound by the viral polymerase L and encapsidated by the viral nucleoprotein N ***(3)***. The expression of the nanoluciferase gene by the VLP-borne L polymerase allows the quantification of tc-VLPs. We previously showed that these tc-VLPs can be neutralized by CCHF-specific antisera ***(4)***, as attested by a decreased nanoluciferase activity when the tc-VLPs were pre-incubated with such sera.

Here, we describe a BSL-2 optimized and down-scaled micro-neutralization assay that allows the detection of CCHF-specific neutralizing antibodies in a 96-well plate setting, both in sera and in libraries of purified monoclonal antibodies. The read-out is a simple luciferase assay and the procedure takes 3 days (Figure 1).

**Figure 1.**
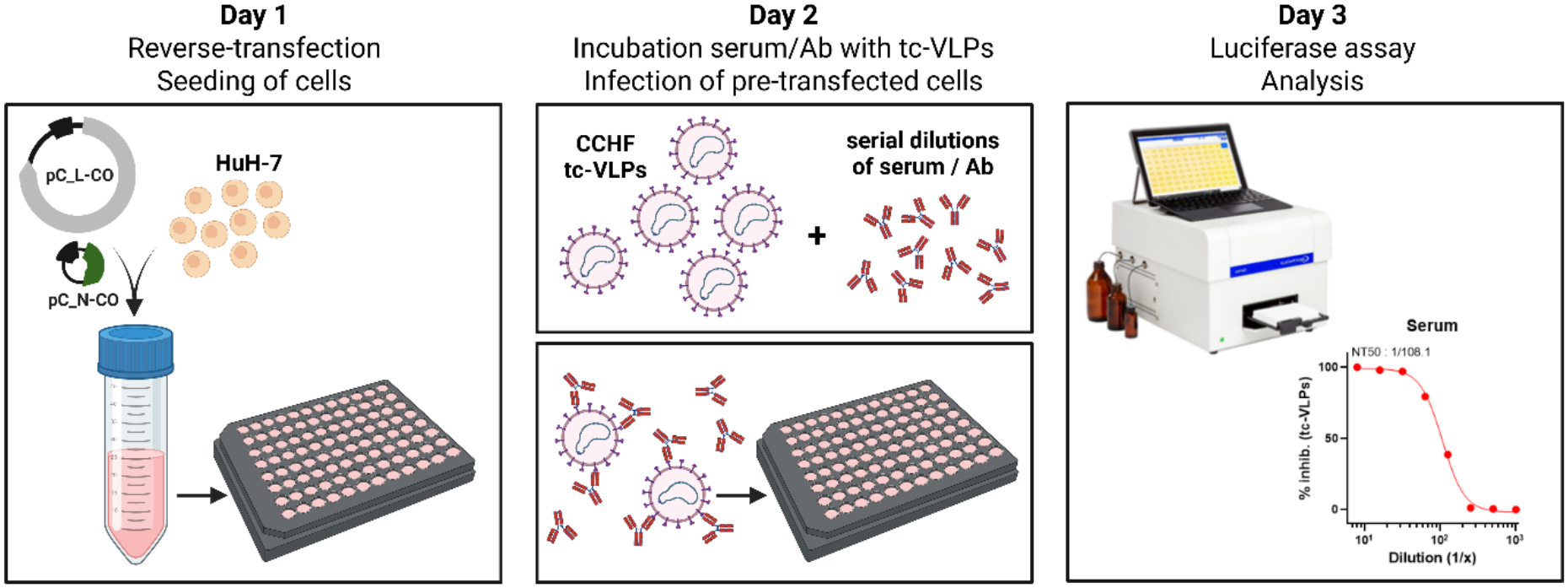
General procedure of the CCHF tc-VLP micro-neutralization assay. Figure created in Biorender.

Day 1: Before being infected with tc-VLPs, HuH-7 cells are pre-transfected with codon-optimized plasmids encoding for CCHFV polymerase L and nucleoprotein N. These two recombinant virus proteins will enhance the transcription and replication of the luciferase-minigenome carried by the incoming nucleocapsids. The cells are seeded in 96-well plates at the same time as the reverse-transfection is done.

Day 2: The sera/antibodies of interest are serial diluted and incubated with CCHF tc-VLPs. The mixes are then used to infect the pre-transfected HuH-7 cells.

Day 3: The cells are lysed, incubated with a luciferase activity detection reagent, and luciferase activity is measured with a luminometer. The determination of neutralization titers is done by a non-linear regression analysis (4-parameters) analysis.

## 2 Materials

### 2.1 Cell culture

1. HuH-7 cells (Human hepatocytes; Cytion)
2. Phosphate Buffer Saline (PBS), calcium-free and magnesium-free
3. Trypsin-EDTA
4. Complete medium: DMEM (Gibco, 41966-029) + 10% heat-inactivated FBS (Gibco)
5. Sterile black 96-well plate with transparent bottom (ie, Greiner 655090, or Corning CLS3603-48EA) (*see* **Note 1**)
6. Reservoirs for multichannel pipetting
7. Multichannel pipet for 100-200μL

### 2.2. Plasmids and DNA transfection

1. plasmid pCAGGS_V5_CCHFV-IbAr_L-CO, in short pC_L-CO (GenBank PZ941510; encodes the CCHFV wild-type polymerase, from a codon-optimized sequence) (*see* **Note 2**)
2. plasmid pCAGGS_CCHFV-IbAr_N-CO, in short pC_N-CO (GenBank PZ941511; encodes the CCHFV nucleocapsid protein, from a codon-optimized sequence) (*see* **Note 2**)
3. TransIT®-LT1 Transfection Reagent (Mirus-Bio, MIR2300)
4. Opti-MEM (ThermoFisher Sci., 31985-062)

### 2.3. Neutralization assay

1. 96 well plates with round bottom (Sarstedt 82.1582.001)
2. DMEM (Gibco, 41966-029)
3. Nunc™ Seals transparent polyester, sterile for cell culture (ThermoFisher Sci., 236366)
4. CCHF tc-VLPs encoding a nanoluciferase reporter gene (*see* **Note 3**)
5. A positive control serum / monoclonal antibody (mAb), a negative control serum, and the sera/mAbs to be tested
6. Orbital shaker that fits into the 37°C, 5% CO_2_ incubator
7. Reservoirs for multichannel pipetting
8. Multichannel pipet for 100-200μL and 10-50 μL

### 2.4. Luciferase assay

1. Nunc™ Seals black vinyl (ThermoFisher Sci., 236703)
2. Reservoirs for multichannel pipetting
3. Multichannel pipet for 100-200μL and 10-50 μL
4. Passive lysis buffer (Promega)
5. Nano-Glo® Luciferase Assay System (Promega)
6. Luminometer, if possible with injectors (e.g. Centro XS microplate luminometer, Berthold Technologies)
7. Software such as Prism from GraphPad, to do the 4-parameter analysis

## 3. Methods

### 3.1. Day 1: Reverse-transfection and seeding of cells

1. General considerations. All work is done under a Class II Biosafety cabinet. The assay is done in a black 96-well plate with transparent bottom. In a single well, there is a final volume of 150 μL: 10,000 HuH-7 cells in 100 μL medium, and 100 ng pC_L-CO and 30 ng pC_N-CO in 50 μL Opti-MEM / transfection reagent. The ratio for the transfection reagent is 2 μg TransIT-LT1 per 1 μg plasmid, in an Opti-MEM solution. All volumes described afterwards are for a master mix of 100 wells, for an assay in one 96-well plate. The conditions “mock” and “tc-VLPs only” should be included in every single plate, even if several plates are required in one experiment (*see* **Note 4**).
2. Preparation of the cells. Remove the medium from the culture flask of HuH-7 cells (*see* **Note 5**). Wash delicately the monolayer once with PBS (*see* **Note 6**) and trypsinize for 3 minutes at 37°C (*see* **Note 7**). Stop the trypsin action by adding DMEM medium with 10% FCS (= complete medium), and centrifuge the cells 5 min at 400 x g. Resuspend the cells in complete medium and count them. Do another centrifugation 5 min at 400 x g, and resuspend the pellet in complete medium, at a concentration of 1×10^6^ cells / mL.
3. Preparation of the plasmid transfection mix. The TransIT-LT1 transfection reagent should be put at room-temperature before use. All plasmid working stocks are at an initial concentration of 100 ng / μL (*see* **Note 8**). For 1 well of a 96-well plate, the transfection is made with 100 ng pC_L-CO and 30 ng pC_N-CO, in 50 μL per well (volume of plasmids + transfection reagent + Opti-MEM). Master-mix for 100 wells: 100 μL pC_L-CO + 30 μL pC_N-CO + 4 844 μL Opti-MEM, in a 50 mL tube (*see* **Note 9**). Tap gently on the side of the tube to mix. Add 26 μL of TransIT-LT1 transfection reagent, and tap gently on the side of the tube to mix. Incubate 20 min at room-temperature.
4. Preparation of the plasmid / cell mix, and seeding in 96-well plate. In the 50 mL tube containing the transfection mix, add 1 mL of the 1×10^6^ cells/mL suspension, and 9 mL of complete medium. Pipet up-and-down slowly with a 10 mL pipet, and transfer into a sterile reservoir. Using a multichannel pipet, distribute 150 μL of the cells / transfection mix per well of a black 96-well plate with transparent bottom. Incubate 20 hours in a 37°C, 5% CO_2_ incubator.

### 3.2. Day 2: Incubation serum/antibodies with CCHF tc-VLPs, and infection of the pre-transfected cells

1. General considerations. All work is done under a Class II Biosafety cabinet. For one 96-well plate, 5 mL of tc-VLPs are needed. Each plate should contain 8 wells with DMEM only (“mock”) and 8 wells with “tc-VLPs only”. On top of these controls, each plate can hold up to 5 different sera/mAbs in duplicate. All calculations are made to be able to infect the cells in duplicate. See Figure 2 for a general outline of the dilutions and a suggestion of a final plate map.
2a. Serial dilution of sera: They are prepared in 2-fold serial dilutions for a final range of 1:8 – 1:1024 when mixed with the tc-VLPs. Sera are first heat-inactivated for 30 min at 56°C. Then all steps are done in a round-bottom 96-well plate, to facilitate handling. According to Figure 2A and for each serum, distribute 20 μL of DMEM in 7 wells. Vortex each heat-inactivated serum, and pipet 20 μL in the first well (that already contains 20 μL DMEM). Proceed to the serial dilution, by pipetting several times up-and-down, and transferring 20 μL from one well to the next. Change the pipet tip between each dilution.
2b. Serial dilution of monoclonal antibodies (mAbs): They are prepared in 3-fold serial dilutions for a final range of 12.5 to 0.006 μg/mL when mixed with the tc-VLPs. First, mAbs are diluted in DMEM to a final concentration of 100 μg/mL. Then all steps are done in a round-bottom 96-well plate, to facilitate handling. For each mAb, distribute 20 μL of DMEM in 7 wells. Pipet 10 μL of mAb (100 μg/mL) in the first well (that already contains 20 μL DMEM). Proceed to the serial dilution, by pipetting several times up-and-down, and transferring 10 μL from one tube to the next. Change the pipet tip between each dilution.
3. Incubation of sera/mAbs with CCHF tc-VLPs. For the controls (Figure 2A): In another round-bottom 96-well plate, distribute 120 μL DMEM in 4 wells, to obtain the condition “mock”. Distribute 105 μL tc-VLPs in another 4 wells (*see* **Note 10**) and add 15 μL of DMEM, to obtain the conditions “tc-VLPs only”. For the sera/mAbs: in the same round-bottom 96-well plate as the controls, distribute 105 μL of tc-VLPs. From the wells with serial dilutions of serum or mAb, transfer 15 μL into each well containing tc-VLPs. Close the wells using a Nunc™ Seal transparent polyester sticker, and incubate for 1 hour at 100 rpm, on a shaker placed inside the 37°C, 5% CO_2_ incubator.
4. Infection of the pre-transfected cells. Using a multichannel pipet, remove the medium from the pre-transfected cells prepared on Day 1, and do one delicate wash with 100 μL PBS. Remove the PBS and add 50 μL of the mixes from the plate “sera/mAbs with CCHF tc-VLPs”, done in the previous step. Change tips and repeat to get duplicates. Incubate for 1 hour in the 37°C, 5% CO_2_ incubator. Then add 100 μL of complete medium to stop the infection (*see* **Note 11**), and incubate for 20 hours in the 37°C, 5% CO_2_ incubator. An example of final plate map is shown in Figure 2B.

**Figure 2.**
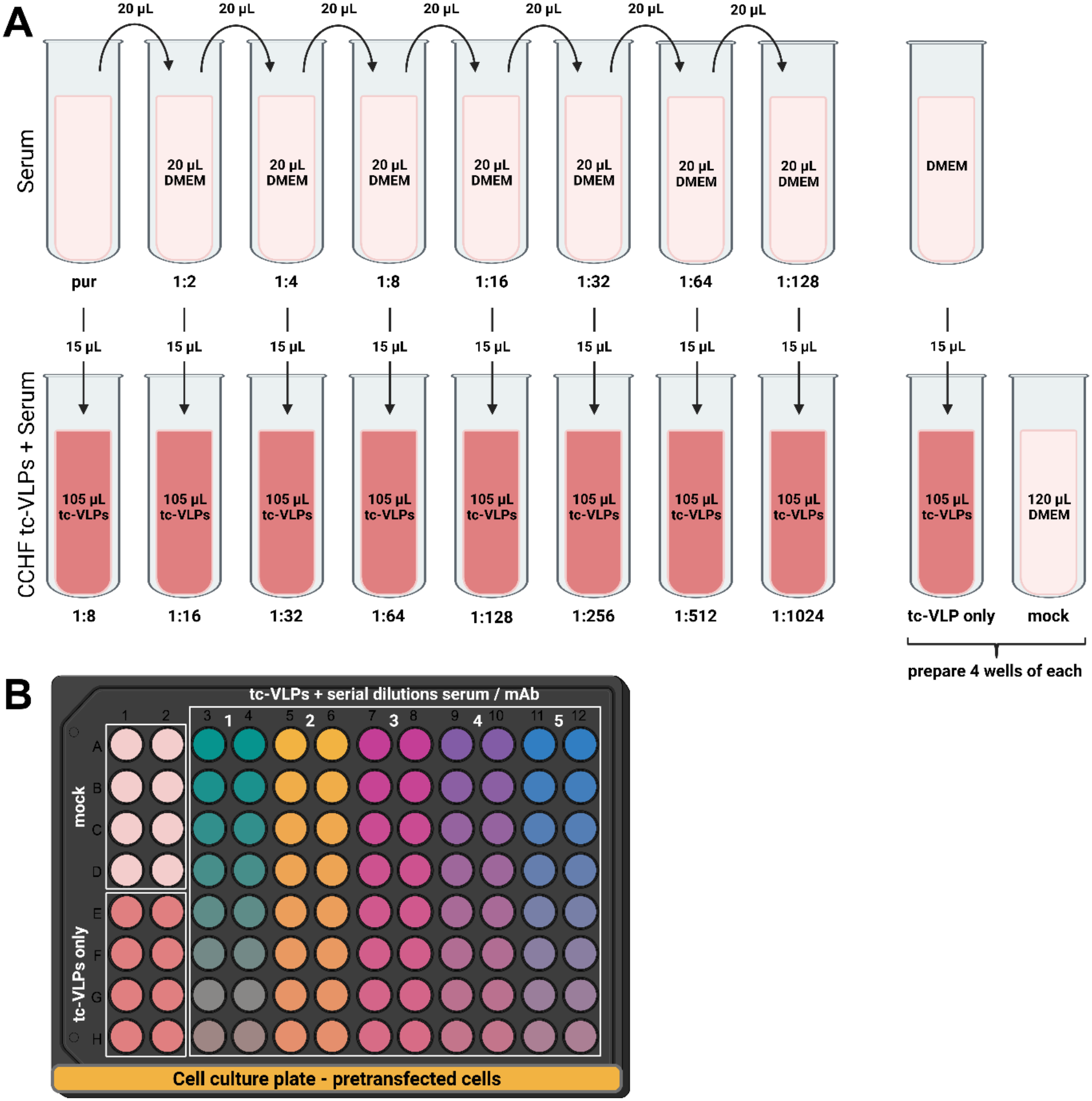
Experimental procedure. A) General outline of the serial dilutions for sera. Volumes are made to be used in duplicate on cells. For the controls “tc-VLP only” and “mock”, prepare 4 wells as described, to do 8-plicates on cells. B) Example of a final plate map, which should always contain 8 wells of “mock”, 8 wells of “tc-VLPs only”, on top of the serum/mAb serial dilutions. Figure created in Biorender.

### 3.3. Day 3: Luciferase assay and analysis

1. General considerations. Follow the manufacturer protocol for the preparation of the Nano-Glo® Luciferase Assay Reagent. A total of 4 mL of reagent is needed per plate.
2. Lysis of the cells. Remove the medium from the wells, and flip the plate on a paper-towel. Stick the black vinyl Nunc seal on the back side of the plate, and make sure all well bottoms are covered (*see* **Note 12**). Using a multichannel pipet, distribute 15 μL of passive lysis buffer 1X (PLB) per well (*see* **Note 13**). Gently swirl the plates 15 min at 150 rpm at room temperature.
3. Luciferase assay. Put the plate in the luminometer with injector system (*see* **Note 14**), and use the following program: inject 40 μL of the Nano-Glo® Luciferase Assay Reagent in all the wells, shake 1 sec, delay 5 min, read luminescence with an integration time of 1 sec.
4. Analysis. Proceed per plate, and refer to the “mock” and “tc-VLP only” values in this plate. For each duplicate, calculate the percentage inhibition using the formula 100*(1-(x-min)/(max-min)), in which “min” is the mean value of the 8 “mock” wells, and “max” is the mean value of the 8 “tc-VLP only” wells. In GraphPad Prism, create a XY table with duplicates in Y, and insert the % reduction values. Perform a non-linear regression analysis, using the setup “[inhibitor] vs response – variable slope (4 parameter)”, without any constrain (*see* **Note 15**). The calculated IC50 is the neutralization titer (NT50) of the serum / mAb. An exemple of analysis curves for positive and negative control sera are shown in Figure 3.

**Figure 3.**
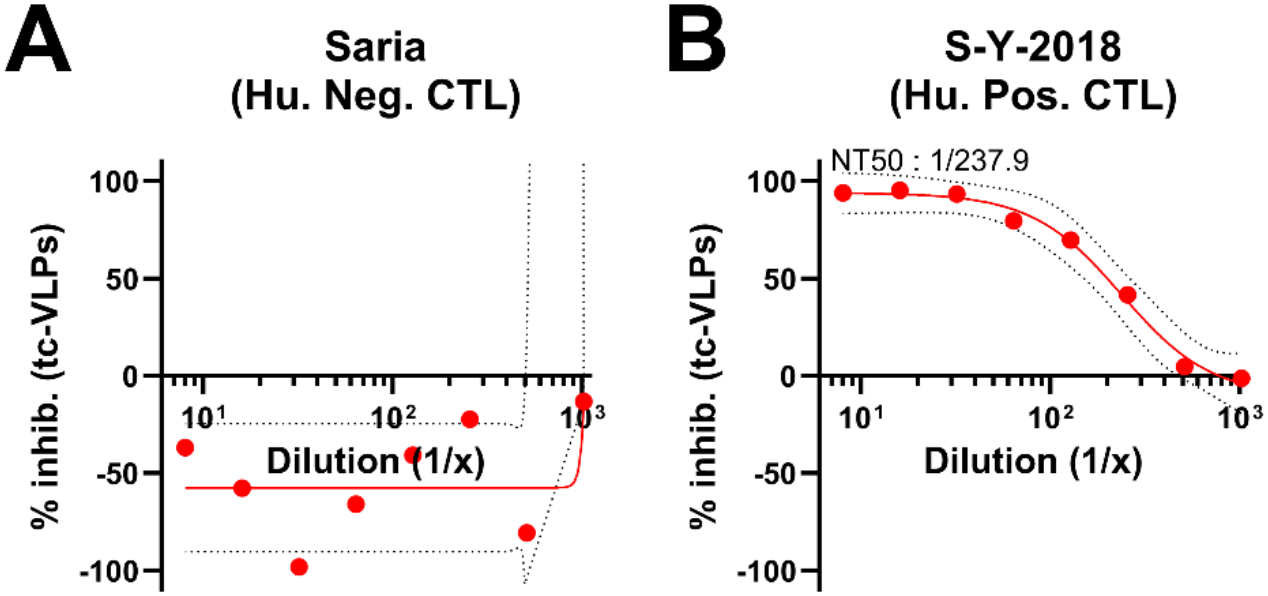
Typical non-linear regression analysis curves for a negative human serum (A) and a positive human serum (B), shown in red. The confidence interval at 95% (IC95%) is shown by the dotted black lines. Mean of 3 independent experiments, done in technical duplicate.

## 4 Notes

1. Although it is generally advised to use white plates for the luciferase assay, comparative experiments done in our lab showed a better signal-to-background ratio when using black plates.
2. Alternatively, the plasmids pCAGGS_V5_CCHFV-IbAr_L and pCAGGS_CCHFV-IbAr_N can be used ***(5)***.
3. The CCHF tc-VLPs harboring a nanoluciferase reporter gene can be produced according to Devignot *et al*.***(3)***.
4. The “mock” and “tc-VLP only” conditions are setting up the background level and the highest RLU (relative light unit), which are needed for the % reduction calculation. These values can vary depending on the time of incubation with the various reagent of the luciferase assay (passive lysis buffer, detection reagent, …) and should hence be included in every plate.
5. The cells should be mycoplasma-free and regularly tested for contaminations. Also, HuH-7 cells should be under-confluent, big and round, without any vacuoles. If they are star-looking, shrunk, or with many small vacuoles, they should not be used for the experiment.
6. PBS should be calcium-free and magnesium-free, to help cell dissociation.
7. Trypsinizing for a longer time can result in aggregation of the HuH-7 cells into white aggregates, which are difficult to resuspend. These aggregates should not be used in the experiment. It is better to trypsinize for a shorter time, and tap the flask on the side to help the cells to detach.
8. The pC_L-CO plasmid is very large and needs special handling. We are using the NucleoBond Xtra Maxi kit (Macherey-Nagel) to make the plasmid stock, with slow pipetting at the resuspension step. To avoid breaking the plasmid when making the dilution stock for the assay, pipet the plasmid into the Opti-MEM and tap delicately on the side of the tube to mix. Do not vortex.
9. The plasmids will be mixed with Opti-MEM, the transfection reagent, and the cell suspension, reaching a volume of 15mL for one 96-well plate.
10. It is difficult to get an accurate titer of tc-VLPs, because it is based on the efficiency of pre-transfection by the pC_L-CO and pC_N-CO plasmids, and the efficiency of infection. Both TCID50 and IF methods are based on the expression levels of the luciferase minigenome, and titers are most often under-estimated. Also, in relation to this, no correlation could be made between tc-VLP titers and luciferase counts in the cells (unpublished observation). So each individual batch of tc-VLPs should be tested with and without a positive control serum, before doing the actual assay. Generally, tc-VLPs with a luciferase count of 500,000 to 1,000,000 RLU when used in a 96-well plate without serum, lead to acceptable results. All related samples, ie comparison of different mAbs or comparison of serum samples collected on a time-course, should be done with the same set of tc-VLPs, on the same experiment, and if possible in the same plate.
11. As tc-VLPs are not able to make new particles, it is not necessary to remove the inoculum.
12. The luciferase assay is based on light emission, so it is very important to protect wells from external and cross-well light sources to avoid background signal.
13. According to the manufacturer, the Nano-Glo® Luciferase Assay Reagent can be added 1:1 on cells in their culture medium. However, we were able to obtain a better signal when lysing the cells first in passive lysis buffer (PLB), and adapting the volumes of reagent needed.
14. Using an injector assures a better reproducibility and homogeneous results within and between experiments. However, it is possible to handle the assay manually, by adding the Nano-Glo® Luciferase Assay Reagent using a multichannel, incubating 5 minutes in a dark room, and then reading the luminescence in the luminometer.
15. Very often, the luminescence counts are greater than the mean of the 8 “tc-VLP only” values, leading to a percentage reduction that is negative. This means that there is no neutralization. While these values could be set to zero, we prefer to keep the original values, to have a S-shape curve fitting the serum/mAb serial dilution, and allowing a more accurate calculation of the NT50.

## 5. Competing Interests Statement

The authors have no conflicts of interest to declare that are relevant to the content of this chapter. Our work was financed by the CCHFever Network supported by the EU under the Health Cooperation Work Programme (grant agreement no. 260427), the CCHFVaccine consortium supported by the EU Horizon 2020 research and innovation program (grant agreement no. 732732), and the Swedish Research Council (VR no. 2018-05766).

## Notes

### Competing Interest Statement

The authors have declared no competing interest.

